## Supplementary figures S1-5 and tables S1-2 for "Structure, Mechanism and Crystallographic fragment screening of the SARS-CoV-2 NSP13 helicase"

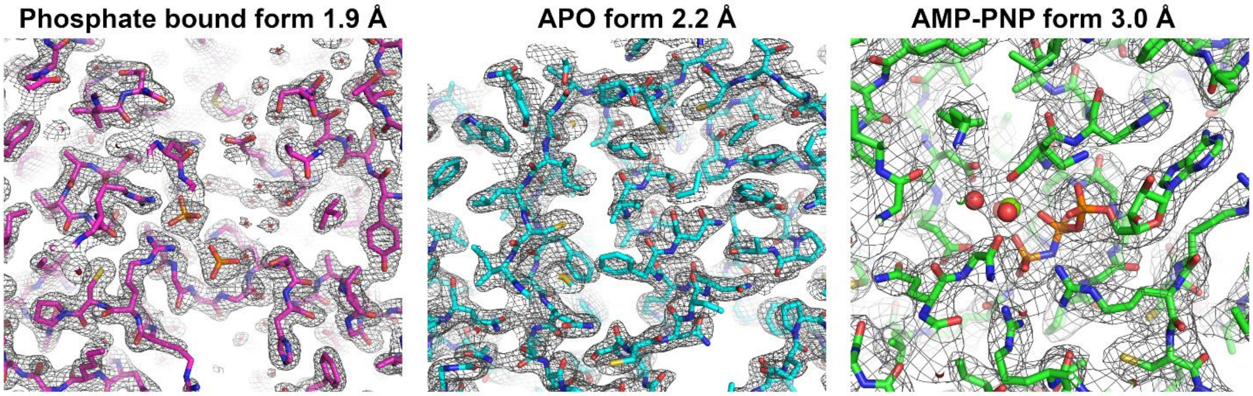


**Figure S1 –** Representative 2$F_o$-1$F_c$ electron density maps of the various NSP13 crystals contoured at 1.2 σ.

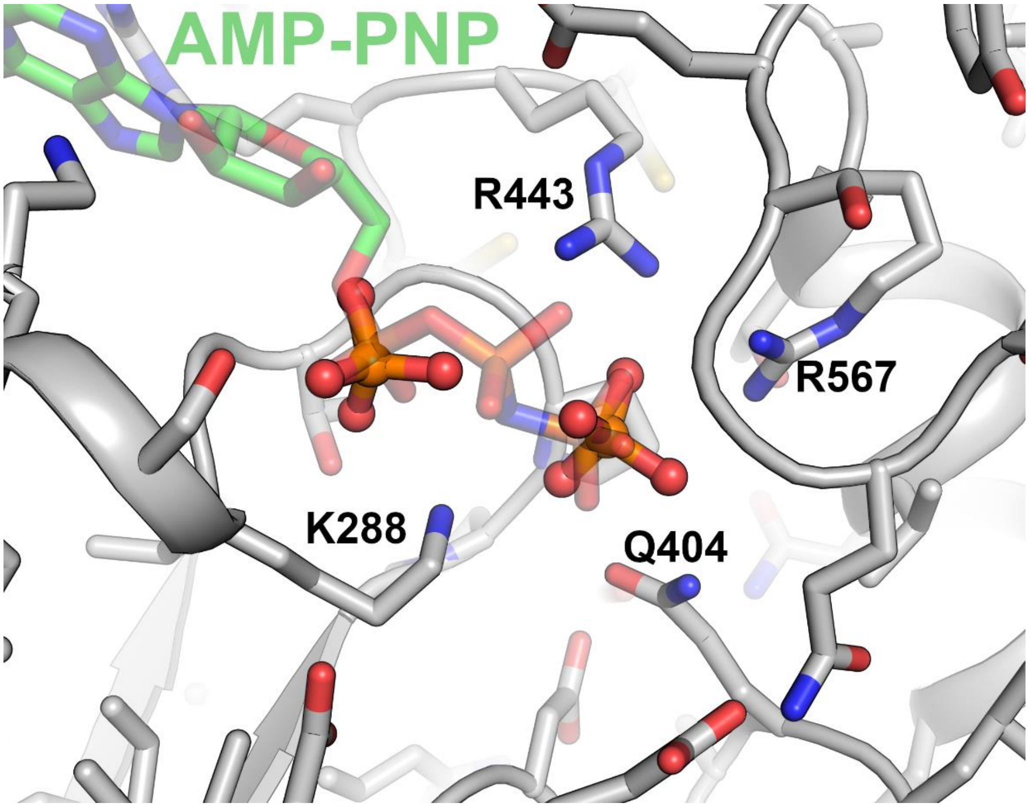


**Figure S2 –** Comparison of the position of the phosphate ions in the phosphate bound crystals with the AMP-PNP in binding mode B. The AMP-PNP is shown in semi-transparent green with the two phosphates in the phosphate bound form occupying positions equivalent to the σ and γ.

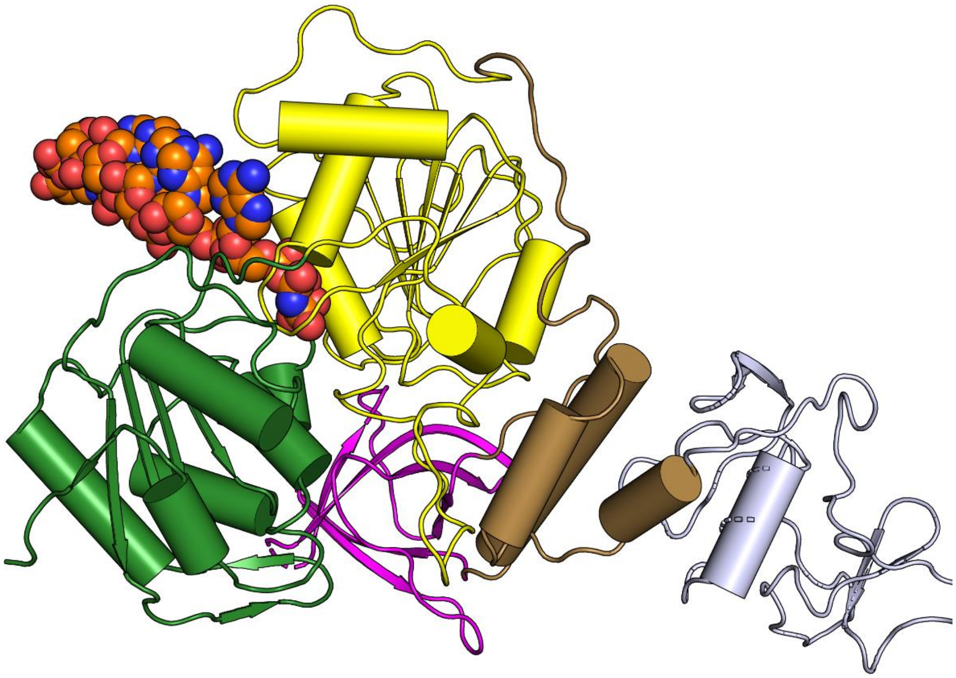

**Figure S3 –** Model of a NSP13 accommodating an RNA containing a 5’ triphosphate into the NSP13 ATPase active site. The model was constructed by connecting a short RNA template with the free 3’ OH of the AMP-PNP nucleotide which points toward the solvent in binding mode A and can be connected without any severe steric clashes.

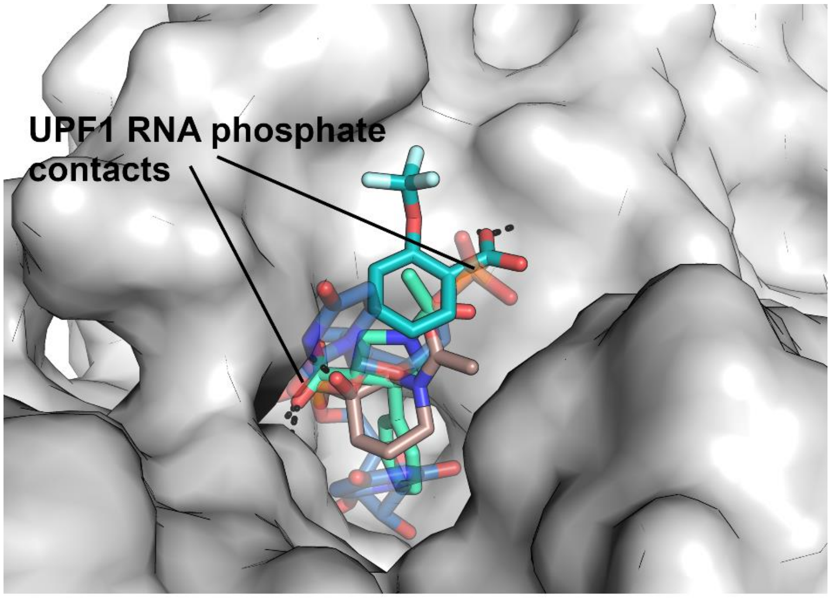


**Figure S4 –** Fragments in the 5' end of the RNA binding channel make contacts to the protein that are shared by conserved RNA phosphate interactions in the UPF-1 RNA structure. The RNA is shown in semi-transparent blue with phosphate positions indicated.

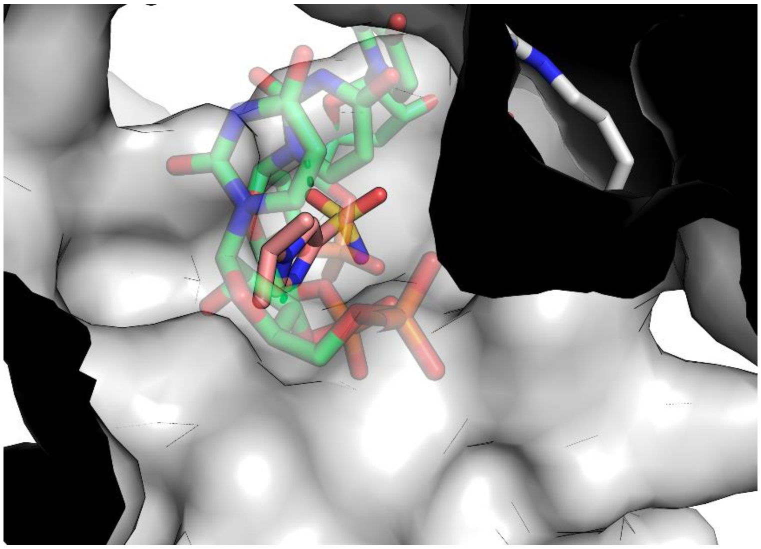

**Figure S5 –** A Fragment in the central RNA binding channel makes contacts to the protein that are shared by conserved RNA phosphate interactions in the UPF-1 RNA structure. The RNA is shown in semi-transparent green.

**Table S1 –** Pairwise RMSD values for overall structure comparisons of individual NSP13 chains. Chains from this study are references by chain ID for the phosphate bound (Phos), APO (APO) and AMP-PNP (PNP) bound forms whilst other structures are referenced by their PDB ID.

| | This study | | | | | | | | SARS–CoV-1 | | Cryo-EM structures | | | | |
|---|---|---|---|---|---|---|---|---|---|---|---|---|---|---|---|
| | Phos-A | Phos-B | APOA-A | APO-B | PNP-A | PNP-B | PNP-C | PNP-D | 6JYT-A | 6JYT-B | 6XEZ-1 | 6XEZ-2 | 7CXM-1 | 7CXM-2 | 7CYQ-1 |
| Phos-A | 0 | 1 | 1.227 | 0.569 | 1.843 | 0.966 | 2.11 | 1.26 | 1.637 | 1.759 | 2.57 | 2.75 | 2.611 | 2.752 | 1.789 |
| Phos-B | | 0 | 0.522 | 1.264 | 2.243 | 0.925 | 1.24 | 0.663 | 1.606 | 1.667 | 2.037 | 2.861 | 2.104 | 2.839 | 1.314 |
| APO-A | | | 0 | 1.331 | 2.406 | 1.02 | 1.05 | 0.913 | 1.654 | 1.638 | 2.287 | 2.805 | 2.335 | 2.871 | 1.562 |
| APO-B | | | | 0 | 1.58 | 0.871 | 1.888 | 1.318 | 1.104 | 1.226 | 2.403 | 2.609 | 2.408 | 2.553 | 1.911 |
| PNP-A | | | | | 0 | 2.17 | 3.05 | 2.41 | 2.578 | 2.575 | 3.112 | 2.072 | 3.002 | 1.649 | 2.435 |
| PNP-B | | | | | | 0 | 1.25 | 0.67 | 1.118 | 1.246 | 2.028 | 2.986 | 2.135 | 3.019 | 1.895 |
| PNP-C | | | | | | | 0 | 0.75 | 1.5 | 1.536 | 2.006 | 3.327 | 2.141 | 3.46 | 1.92 |
| PNP-D | | | | | | | | 0 | 1.141 | 1.189 | 1.857 | 3.007 | 1.996 | 3.083 | 1.566 |
| 6JYT-A | | | | | | | | | 0 | 0.382 | 1.838 | 3.334 | 1.969 | 3.399 | 2.156 |
| 6JYT-B | | | | | | | | | | 0 | 1.717 | 3.077 | 1.901 | 3.232 | 2.156 |
| 6XEZ-1 | | | | | | | | | | | 0 | 2.849 | 0.866 | 3.227 | 2.24 |
| 6XEZ-2 | | | | | | | | | | | | 0 | 2.744 | 0.843 | 2.768 |
| 7CXM-1 | | | | | | | | | | | | | 0 | 2.878 | 2.18 |
| 7CXM-2 | | | | | | | | | | | | | | 0 | 2.742 |
| 7CYQ-1 | | | | | | | | | | | | | | | 0 |

**Table S2**: Structures and binding sites of NSP13 fragment hits

| PDBID | Ligand | Binding Location | Binding Pocket | Resolution (Å) |
|---|---|---|---|---|
| 5RL7 | 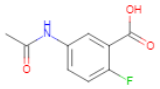 Z364321922 | 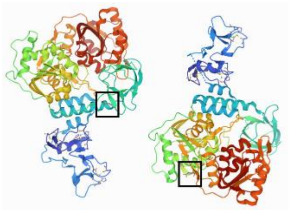 Nucleotide pocket A & RNA 3' B | 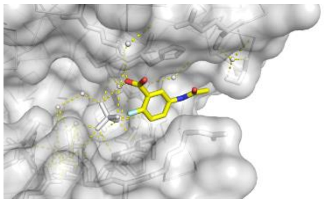 Nucleotide pocket A | 1.89 |
| 5RLV | 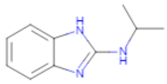 Z2467208649 | 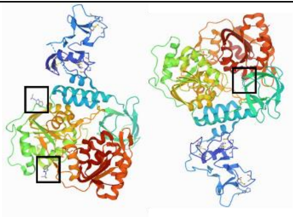 Nucleotide pocket A & Other A & RNA 5' proximal B | 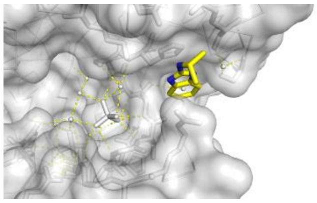 Nucleotide pocket A | 2.21 |
| 5RLY | 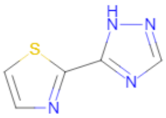 Z2027049478 | 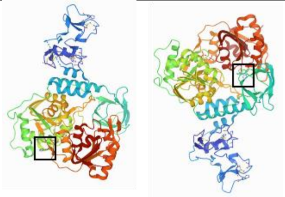 Nucleotide pocket A & RNA 5' proximal B | 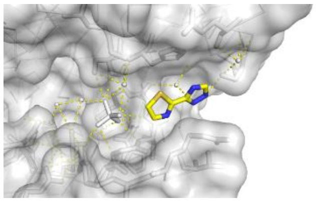 Nucleotide pocket A | 2.43 |
| 5RLS | 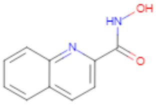 Z59181945 | 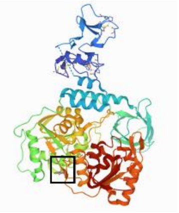 Nucleotide pocket A | 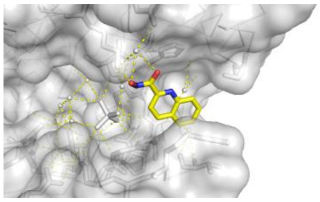 Nucleotide pocket A | 2.28 |
| 5RLN | 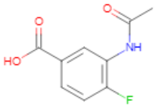 Z364328788 | 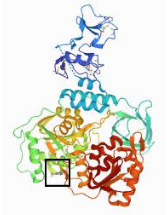 Nucleotide pocket A | 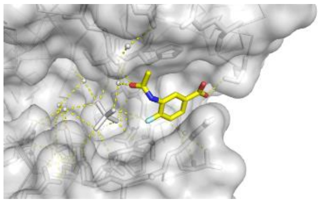 Nucleotide pocket A | 2.15 |
| 7NNG | 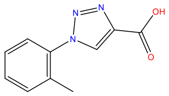 Z2327226104 | 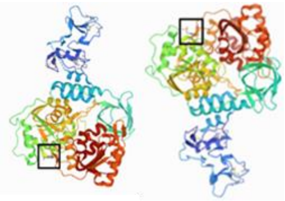 Nucleotide pocket A & B | 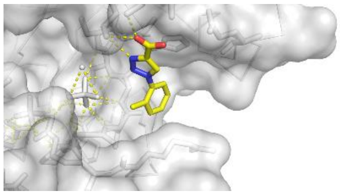 Nucleotide pocket A | 2.38 |

| | | | | |
|---|---|---|---|---|
| 5RL9 | 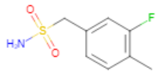 Z1703168683 | 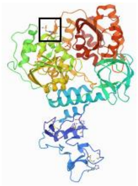 Nucleotide pocket B | 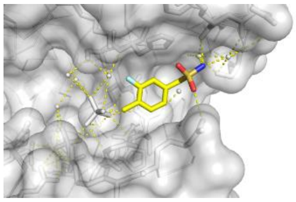 Nucleotide pocket B | 1.79 |
| 5RLI | 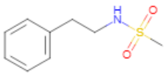 Z45617795 | 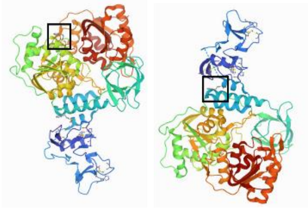 Nucleotide pocket B & Stalk A | 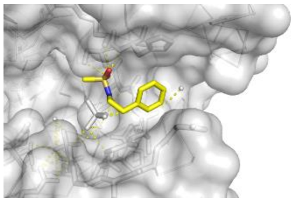 Nucleotide pocket B | 2.26 |
| 5RLJ | 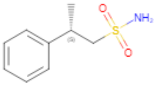 Z1407673036 | 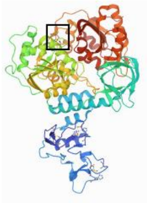 Nucleotide pocket B | 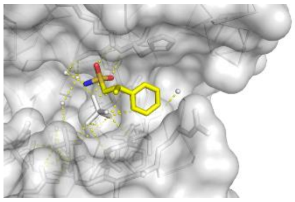 Nucleotide pocket B | 1.88 |
| 5RLO | 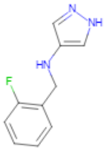 Z1454310449 | 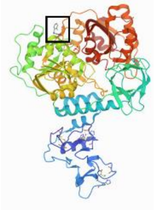 Nucleotide pocket B | 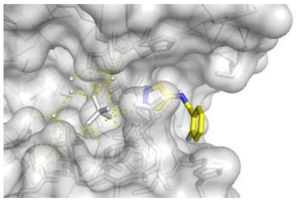 Nucleotide pocket B | 2.10 |
| 5RLR | 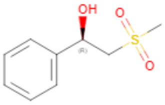 Z822382694 | 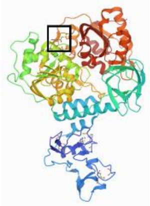 Nucleotide pocket B | 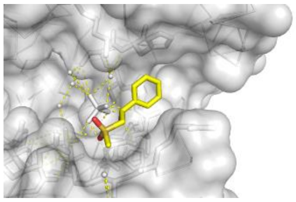 Nucleotide pocket B | 2.32 |
| 5RLW | 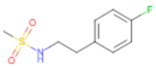 Z45705015 | 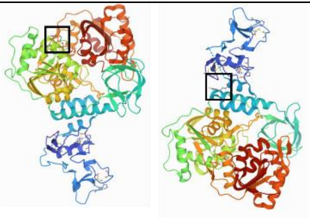 Nucleotide pocket B & Stalk A | 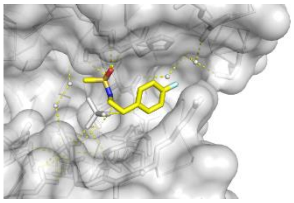 Nucleotide pocket B | 1.97 |

| | | | | |
|---|---|---|---|---|
| 5RM2 | 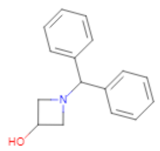 Z1741964527 | 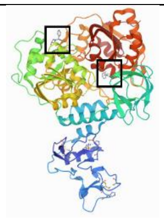 Nucleotide pocket B & RNA 5' Proximal B | 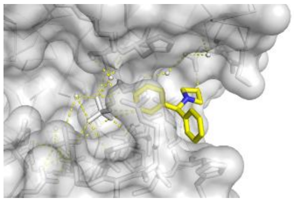 Nucleotide pocket B | 1.82 |
| 5RM7 | 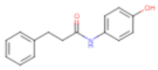 Z69118333 | 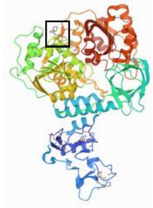 Nucleotide pocket B | 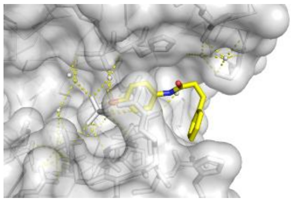 Nucleotide pocket B | 1.84 |
| 5RLL | 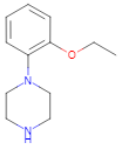 Z425387594 | 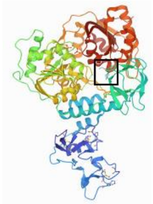 RNA 5' Proximal B | 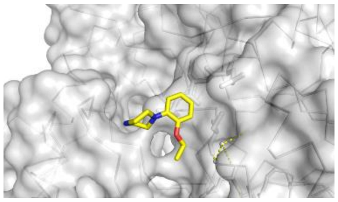 RNA 5' Proximal B | 2.08 |
| 5RLE | 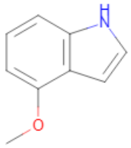 Z1429867185 | 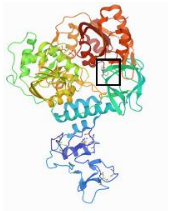 RNA 5' Proximal B | 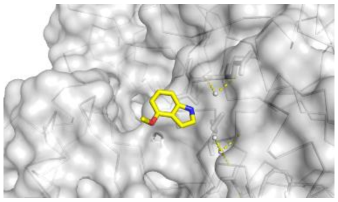 RNA 5' Proximal B | 2.27 |
| 5RLP | 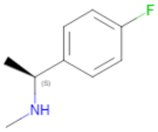 Z166605480 | 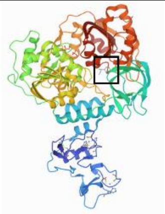 RNA 5' Proximal B | 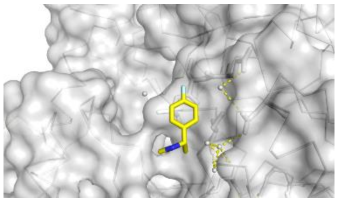 RNA 5' Proximal B | 2.56 |
| 5RMK | 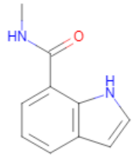 Z1273312153 | 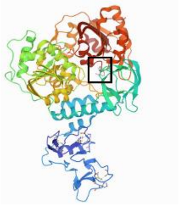 RNA 5' Proximal B | 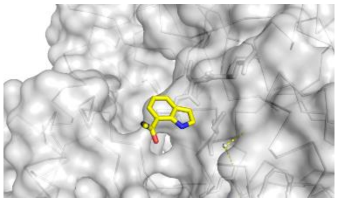 RNA 5' Proximal B | 2.08 |

| | | | | |
|---|---|---|---|---|
| 5RLH | 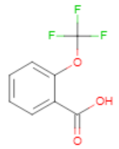 Z2856434778 | 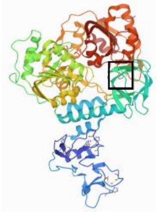 RNA 5’ B | 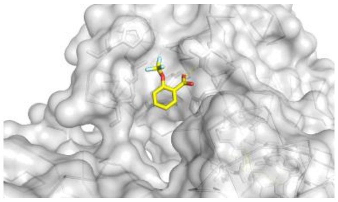 RNA 5’ B | 2.38 |
| 5RMM | 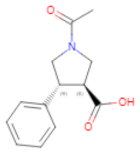 POB0066 | 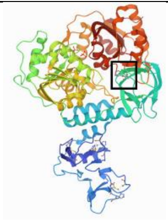 RNA 5’ B | 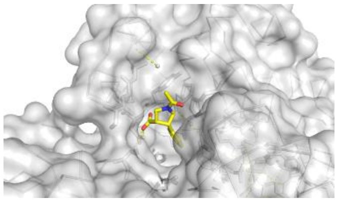 RNA 5’ B | 2.20 |
| 5RLZ | 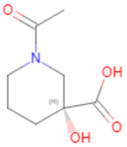 Z2293643386 | 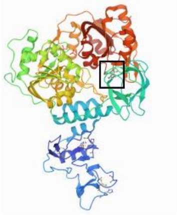 RNA 5’ B | 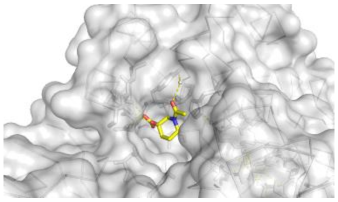 RNA 5’ B | 1.97 |
| 5RL6 | 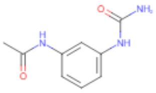 Z198195770 | 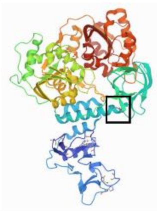 RNA 3’ B | 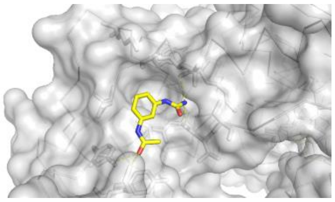 RNA 3’ B | 1.92 |
| 5RLU | 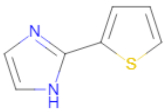 Z744754722 | 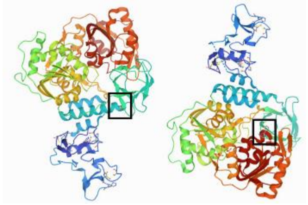 RNA 3’ B & Other A | 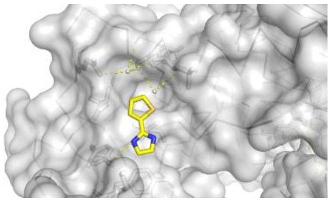 RNA 3’ B | 2.35 |
| 5RL8 | 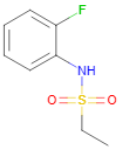 Z53825177 | 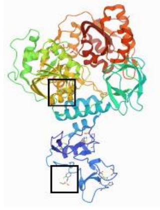 RNA 3’ B & Zinc B | 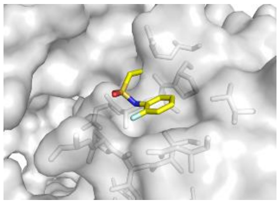 RNA 3’ B | 2.21 |

| | | | | |
|---|---|---|---|---|
| 5RMC | 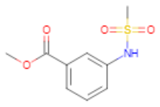 Z24758179 | 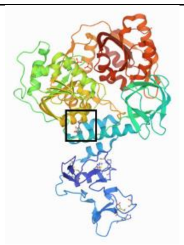 RNA 3’ B | 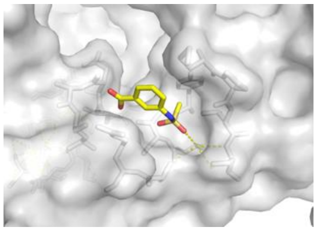 RNA 3’ B | 2.15 |
| 5RLK | 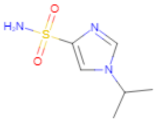 Z1509882419 | 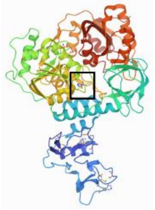 RNA central B | 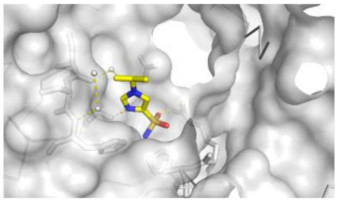 RNA central B | 1.96 |
| 5RML | 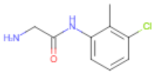 Z85956652 | 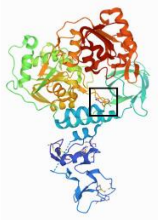 RNA central A | 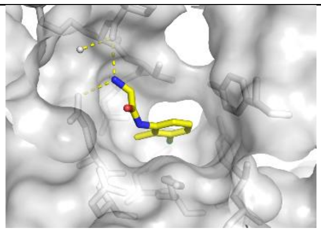 RNA central A | 2.43 |
| 5RLB | 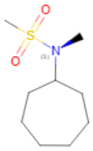 Z216450634 | 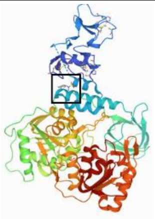 Stalk A | 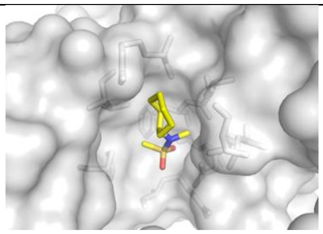 Stalk A | 1.98 |
| 5RMD | 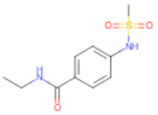 Z57614330 | 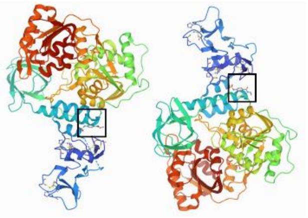 Stalk A & Stalk B | 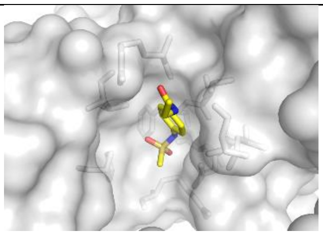 Stalk A | 1.92 |
| 5RLC | 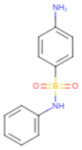 Z56923284 | 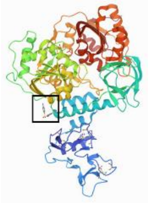 Stalk B | 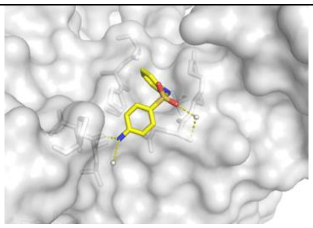 Stalk B | 1.92 |

| | | | | |
|---|---|---|---|---|
| 5RLD | 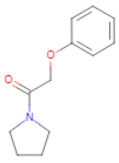 Z19735981 | 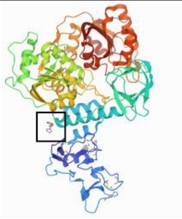 Stalk B | 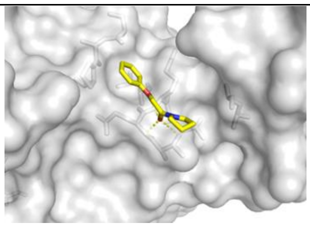 Stalk B | 2.23 |
| 5RM0 | 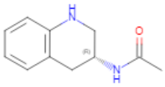 Z1492796719 | 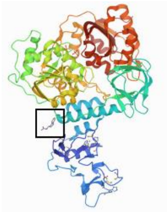 Stalk B | 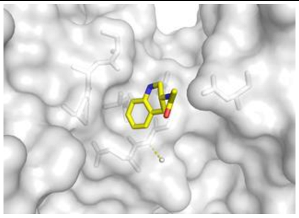 Stalk B | 1.91 |
| 5RM1 | 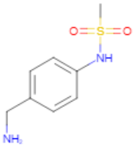 Z426041412 | 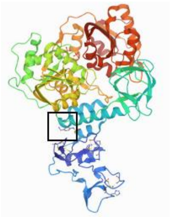 Stalk B | 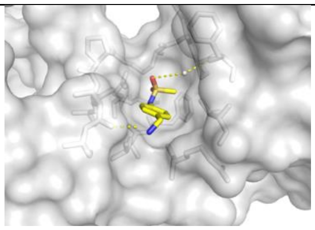 Stalk B | 1.90 |
| 5RME | 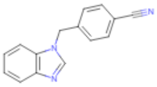 Z26333434 | 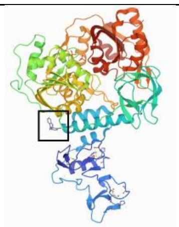 Stalk B | 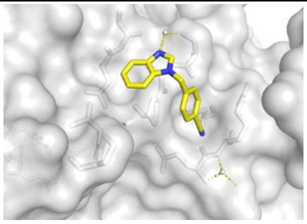 Stalk B | 2.23 |
| 5RLT | 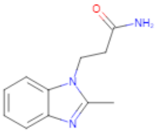 Z53116498 | 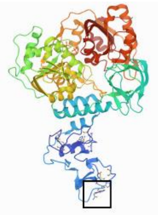 Zinc B | 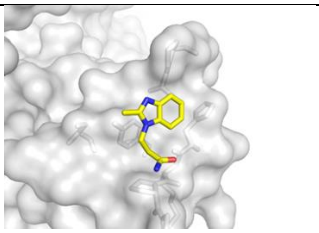 Zinc B | 2.43 |
| 5RLM | 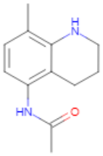 Z1650168321 | 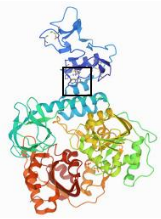 Zinc B | 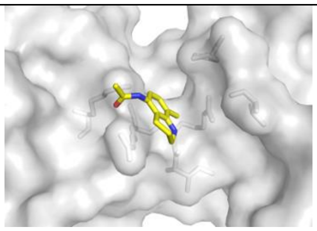 Zinc B | 1.86 |

| | | | | |
|---|---|---|---|---|
| 5RMI | 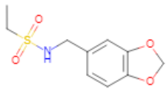<br>Z53860899 | 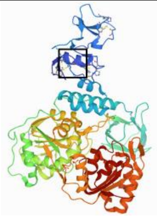<br>Zinc B | 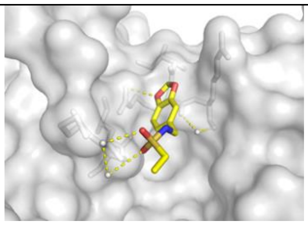<br>Zinc B | 2.12 |
| 5RLF | 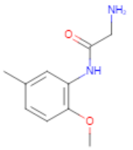<br>Z235341991 | 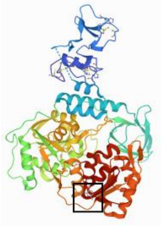<br>2A domain A | 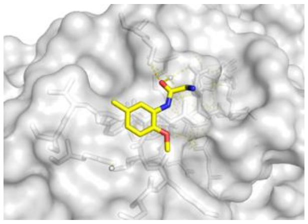<br>2A domain A | 2.23 |
| 5RLQ | 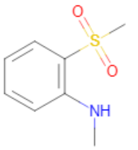<br>Z285782452 | 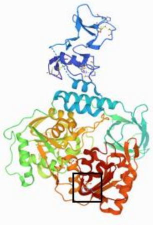<br>2A domain A | 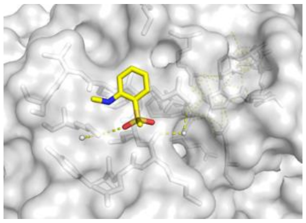<br>2A domain A | 2.23 |
| 5RLG | 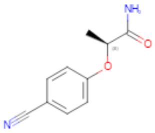<br>Z19739650 | 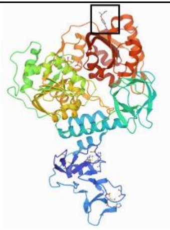<br>2A domain B | 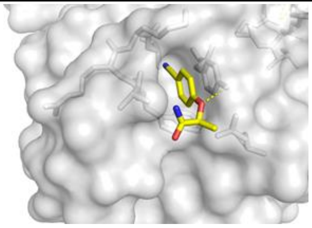<br>2A domain B | 1.96 |
| 5RM3 | 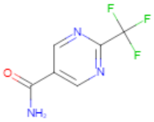<br>Z1745658474 | 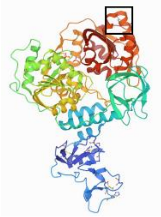<br>C-terminus B | 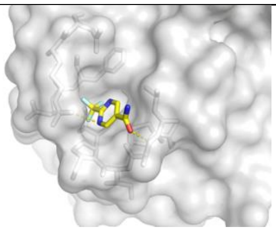<br>C-terminus B | 2.09 |
| 5RM6 | 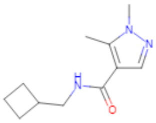<br>Z396380540 | 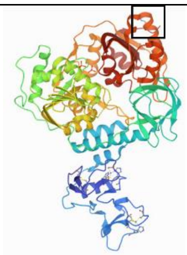<br>C-terminus B | 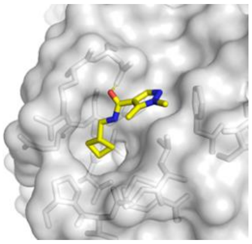<br>C-terminus B | 2.13 |

| | | | | |
|---|---|---|---|---|
| 5RM9 | 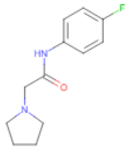 Z2856434942 | 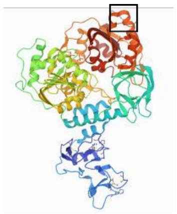 C-terminus B | 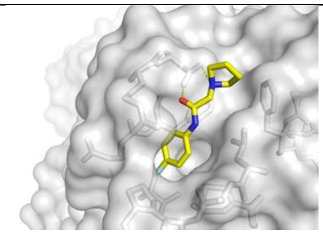 C-terminus B | 2.08 |
| 5RMG | 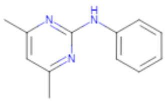 Z285675722 | 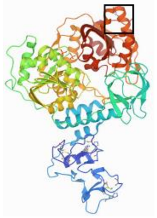 C-terminus B | 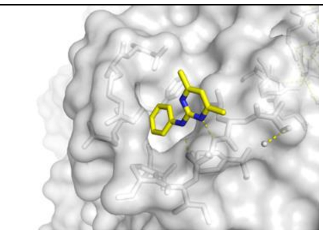 C-terminus B | 2.12 |
| 5RMJ | 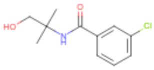 Z68299550 | 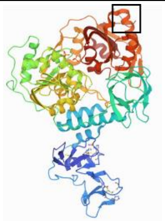 C-terminus B | 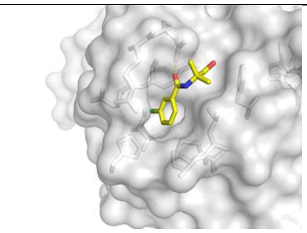 C-terminus B | 2.10 |
| 5RMB | 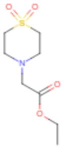 Z2856434920 | 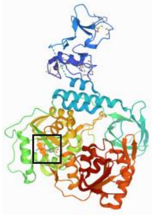 Interface 1A-2A A | 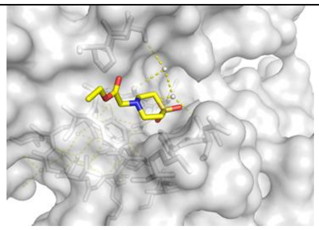 Interface 1A-2A A | 2.21 |
| 5RMF | 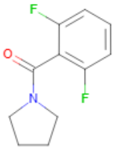 Z54226006 | 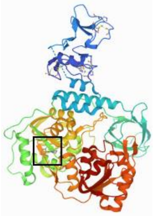 Interface 1A-2A A | 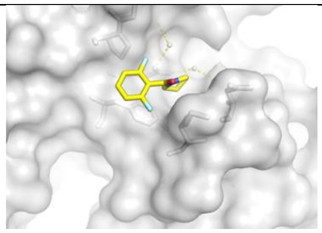 Interface 1A-2A A | 2.23 |
| 5RM4 | 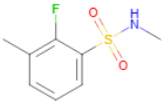 Z1639162606 | 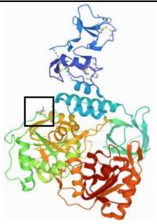 1A domain A | 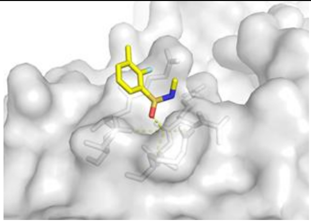 1A domain A | 2.96 |

| | | | | |
|---|---|---|---|---|
| 5RM5 | 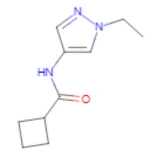 Z373768900 | 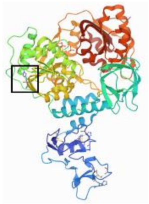 1A domain B | 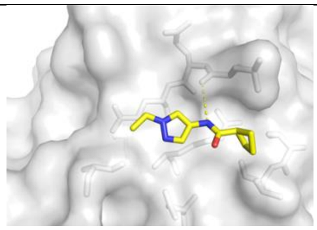 1A domain B | 2.06 |
| 5RM8 | 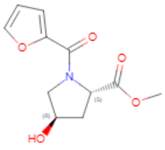 Z1614545742 | 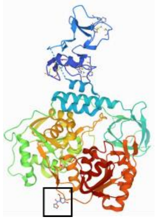 2A domain A | 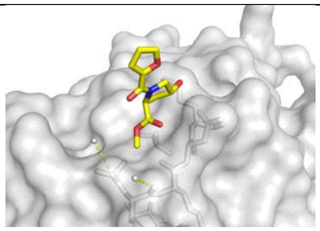 2A domain A | 2.14 |
| 5RMA | 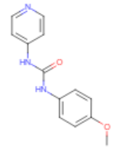 Z321318226 | 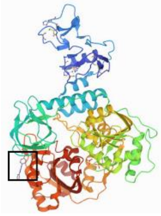 1B-2A interface B | 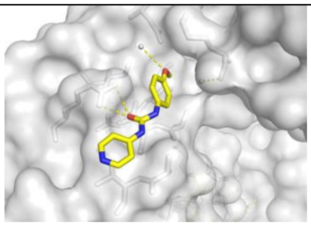 1B-2A interface B | 1.89 |
| 5RMH | 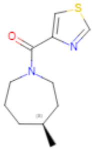 Z1101755952 | 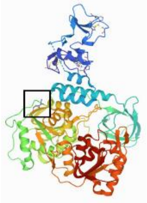 1A domain A | 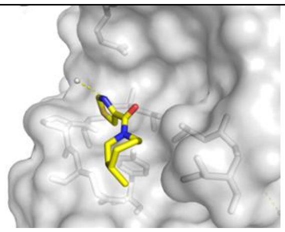 1A domain A | 2.02 |
